# Bespoke single cell molecular and tissue-scale analysis reveals mechanisms underpinning development and disease in complex developing cell populations

**DOI:** 10.1101/2023.10.11.561904

**Authors:** Magdalena E Strauss, Mai-Linh Nu Ton, Samantha Mason, Jaana Bagri, Luke TG Harland, Ivan Imaz-Rosshandler, Nicola K Wilson, Jennifer Nichols, Richard CV Tyser, Berthold Göttgens, John C Marioni, Carolina Guibentif

## Abstract

Perturbation studies using gene knockouts have become a key tool for understanding the roles of regulatory genes in development and disease. Here we systematically characterise the knockout effects of the key developmental regulators *T* and *Mixl1* in chimeric mouse embryos during gastrulation and organogenesis. We present a comprehensive and effective suite of statistical tools for systematic characterisation of effects at the level of differential abundance of cell types, lineage development, and gene dysregulation. Applying our computational approach to a novel chimera data set with *Mixl1* knockout reveals a disruption in Epicardium development in the absence of *Mixl1*, characterized by lack of upregulation of the key transcription factor *Tbx18* and the Wnt regulator *Sfrp5*, and by dysregulation of the recently identified juxta-cardiac field. Finally, we demonstrate the wider utility of our framework by applying it to published acute myeloid leukemia (AML) patient data, and show how different responses to therapy are reflected in changes in gene expression along the myeloid trajectory between healthy and AML patients.

## Background

Using CRISPR knockouts in conjunction with a single-cell transcriptomic readout has become an important tool for understanding gene function. While a number of studies have focused on the development of experimental and analysis techniques for large scale screens, where up to thousands of genes are knocked down in cell lines [1], others have focused on smaller-scale knockout analysis in more complex settings, such as mouse models [2] or organoids [3].

Within the context of such lower-throughput but high information gain model systems, the generation of embryonic chimeras, where mutant cells are injected into wildtype (WT) embryos at the blastocyst stage, is a powerful tool to study the function of essential developmental transcription factors [4–6](Fig. 1A). In the resulting chimeras, WT cells develop normally, establishing signalling gradients necessary for the embryo to develop, while the effects of the knockout can be observed by studying the descendants of the injected mutant cells. Chimeras can therefore reveal cell-autonomous effects of essential gene knockouts. Furthermore, single-cell profiling for chimeras enables the comprehensive study of knockout effects well beyond differences in organ contribution (Fig. 1B, Supp. Fig. 1AB).

**Fig. 1:**
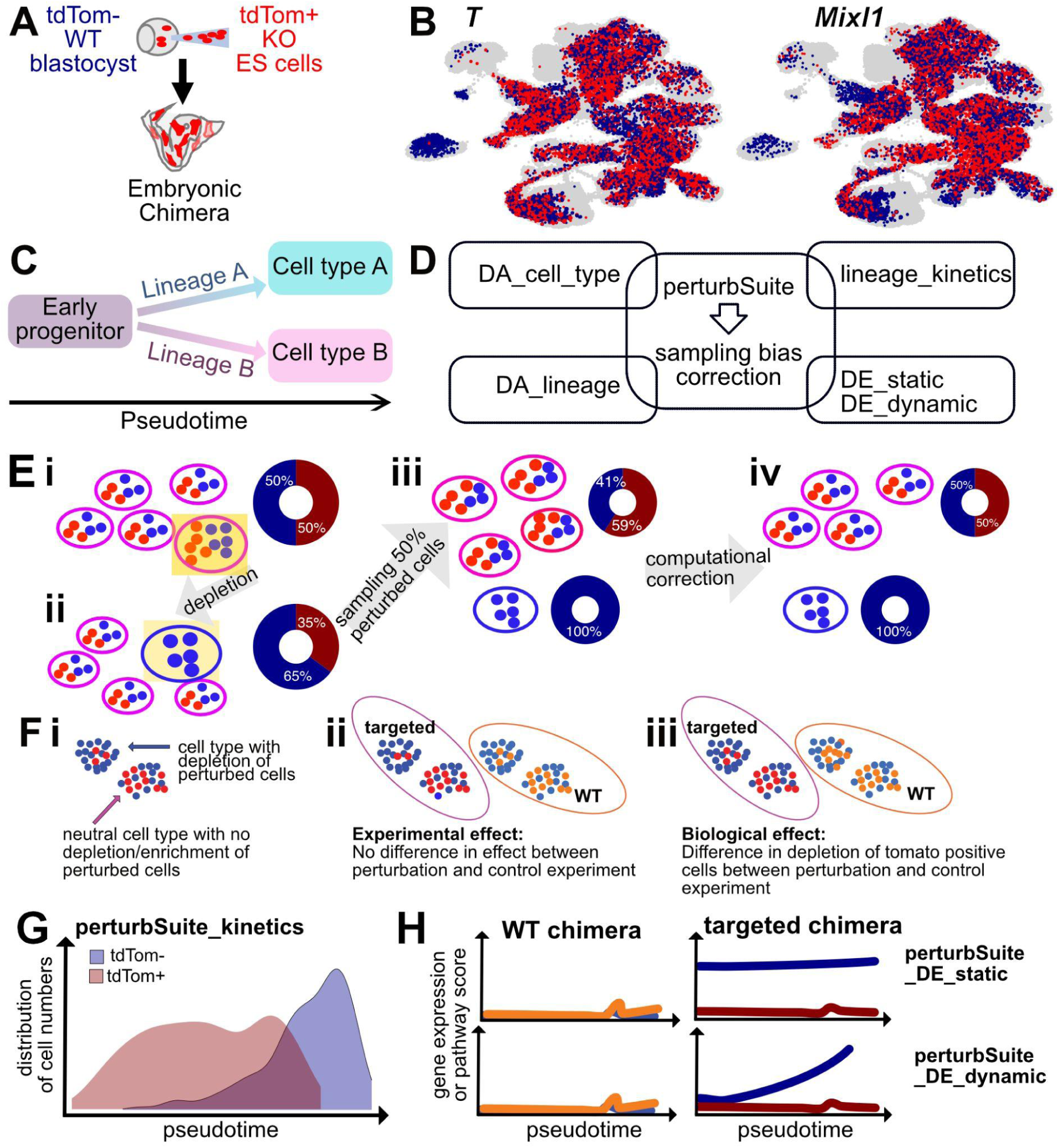
An overview of perturbSuite application to chimera data sets. A) Chimeric embryos were created by injecting mutant tdTom^+^ cells into wildtype tdTom^−^ (WT) embryos at the blastocyst stage. B) Chimeric mouse embryos mapped to the extended mouse gastrulation atlas (reference atlas cells in grey, tdTom^+^ cells in red, tdTom^−^ cells in blue). C) Illustration of lineages. D) Overview of perturbSuite. E) Correction of sampling bias; i) Cell counts from an embryo with knockout cells in red, each cell type represented by a ellipsoid; ii) depletion of one cell type only; iii) this depletion is unknown before data analysis, therefore a fixed proportion of cells with fluorescent markers is sampled, leading to cell types seemingly enriched for knockout cells; (iv) computational correction. F) i) underrepresentation of tdTom^+^ cells (red) for one cell type, ii) experimental effect: similar level of underrepresentation of tdTom^+^ cells for the knockout and WT chimeras, iii) biological effect. G) Illustration of perturbSuite_kinetics. H) Static (top) and dynamic DE (bottom): changes along a trajectory are different for tdTom^+^ versus tdTom^−^ cells of the knockout chimera (right) and this difference is significantly larger than for the WT control experiment (left).

Although powerful, single-cell chimera studies are challenging to interpret, with sampling biases and other technical factors potentially confounding results. In particular, the systematic assessment of the impact of knockouts on lineage progression, on gene expression, and on the interaction of these effects remains challenging. To address these problems, we herein present perturbSuite, a novel set of computational tools specifically designed for the analysis of perturbations in complex developing cell populations, and therefore ideally suited to chimera studies. We validated the proposed analysis framework using chimeric *T/Brachyury* embryos published earlier [5], and then applied it to previously unpublished *Mixl1*^-/-^ chimeric mouse embryo data to reveal - at whole embryo scale - the molecular and tissue-scale consequences of loss of function for this key developmental regulator. Finally, we illustrate the broad utility of our new analysis framework by using it to uncover mechanisms of response to therapy in the context of acute myeloid leukemia (AML) progression.

## Results

### PerturbSuite leverages single-cell profiling to analyse developmental perturbations

We considered two independent chimera data sets, both generated at Embryonic Day (E) 8.5 of mouse development: i) a previously published data set that studied the cell autonomous function of *T* for validation; and ii) a newly generated data set created to study the role of *Mixl1* for new insights into early organogenesis. Additionally, we used data from a control experiment, where wildtype (WT) cells were injected at the blastocyst stage to create control chimeras; this was also profiled at E8.5. In all chimeras, we define the progeny of injected cells as tdTomato positive cells (tdTom^+^, Fig. 1AB), as the injected cells were labelled with the fluorescent marker tdTomato. The wildtype (host) cells lack tdTomato expression and are referred to as tdTomato negative (tdTom^−^).

Profiling the chimeric embryos using single-cell RNA-sequencing (scRNA-seq), in conjunction with reference single-cell atlases of early development, opens exciting possibilities for studying the effects of perturbations during cellular diversification (Fig. 1B, Supp. Fig. 1A-C). For instance, we can explore whether the perturbed cells are depleted or enriched in certain cell types or lineages and assess how far along specific lineages perturbed cells are able to progress (Fig. 1CD). Fig. 1D illustrates the components of our comprehensive perturbSuite framework: comprehensive understanding of knockout effects includes effects on cell type abundance (perturbSuite_DA_cell_type), on lineage abundance (perturbSuite_DA_lineage), on delay along lineage development (perturbSuite_kinetics), and on differential expression (DE, perturbSuite_DE) for a population of cells without considering lineage development (perturbSuite_DE_static) or DE patterns along lineages (perturbSuite_DE_dynamic).

Studying perturbations in complex systems like embryonic chimeras requires attention to specific challenges. First, if a perturbation leads to a strong reduction of some cell types, then the unaffected cell types will appear as if they were enriched in the perturbed cell population. We present an approach to address this bias based on subsampling (Fig. 1E). Second, there are both batch and experimental effects that confound biological signal. To control for both batch and experimental effects, all analyses were framed in terms of two controls; first there is the internal control of unperturbed host cells from the same sample, which is not affected by batch effects. Experimentally, chimera generation may lead to transcriptional differences between the injected tdTom^+^ cells and the tdTom^−^ wildtype host cells even if the injected cells are wildtype (WT) and do not have a knockout. This experimental effect can be corrected for to identify true biological signal (Supp. Fig. 1CD), using the external control, the WT chimeras introduced earlier. However, in contrast to the difference between tdTom+ and tdTom-cells in the same chimera, the comparison between WT and knockout chimeras is usually affected by batch effects. PerturbSuite therefore performs differential abundance (DA) and DE testing using both the internal and external control, thus accounting for both batch and experimental effects (Fig. 1F). This enables the identification of dynamic shifts in lineage development (perturbSuite_kinetics; Fig. 1G) and DE of genes and activity of pathways (Fig. 1H).

In addition to the novel computational approaches, our framework also uses a substantially extended reference atlas of mouse gastrulation[7], which contains four times as many cells with updated cell type annotations, and adds four additional 6-hour timepoints between E8.5 and E9.5 that more completely represents early organogenesis compared to the reference atlas used in previous work [4] (Fig. 1B and Supp. Fig. 1A-C). This allowed us to identify chimeric cells that were similar to cells present in later stage embryos (E8.75 and later, Supp. Fig. 1D).

### PerturbSuite characterises impact of *T* knockout on embryonic development and reveals its role for limb development

We used all elements of perturbSuite (Fig. 1D) to obtain a comprehensive view of the effects of *T* knockout. First, the cell type level analysis of *T*^-/-^ chimeras with perturbSuite_DA_cell_type confirmed previous results [5], with depletion of Somitic Mesoderm, Presomitic Mesoderm, Intermediate Mesoderm and Notochord in the mutant cells and enrichment for NMPs. In addition, we observed a broad depletion effect of other mesodermally-derived lineages including endothelial cell types, namely Embryo proper, Venous and Allantois Endothelium, as well as Mesenchyme, Yolk sac mesothelium, and Allantois (Fig. 2A). We also observed enrichment of Caudal Epiblast cells, which was only present at earlier time points in wildtype embryos, suggesting a developmental delay of mutant cells. Additionally, our pipeline facilitates assessment of time-dependent cellular abundance changes, as seen for example for Blood Progenitors and Cranial Mesoderm, which showed significant changes in relative abundance across different time points (Fig. 2B). We also noted that accounting for the bias illustrated in Fig. 1E is important for avoiding false conclusions about cell types being enriched for knockout cells (Supp. Fig. 2A, which compares perturbSuite_DA with our bias correction to the same method, but with the bias correction left out).

**Fig. 2:**
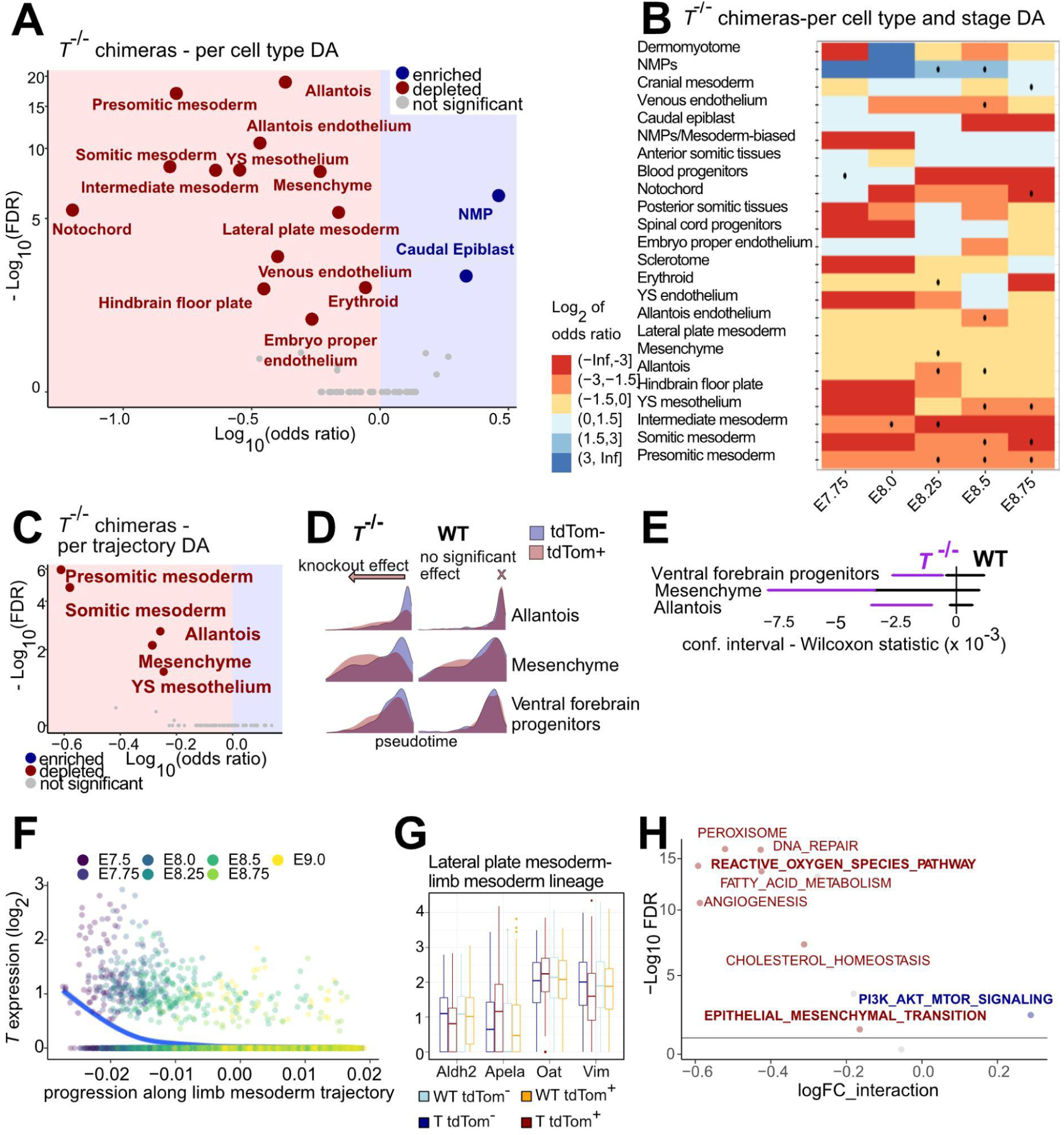
PerturbSuite confirms existing results and reveals mechanistic insights into the role of *T* in the development of limb mesoderm. A) DA at the cell type level for *T^-/-^* chimera knockout cells (perturbSuite_DA_cell_type). B) Per-stage DA at cell type level. Significance at an FDR<0.1 is indicated by a dot. C) DA of Waddington Optimal Transport (WOT) trajectories leading to the listed cell types at E9.25 (perturb_DA_lineage). D) Pseudotime distributions of tdTom^+^ (red) and tdTom^−^ (blue) cells within *T^-/-^* and WT chimeras. E) Confidence interval for location parameter of Wilcoxon test (perturbSuite_kinetics). Confidence intervals for *T*^-/-^ chimeras (purple bars) not overlapping 0 or the confidence intervals for WT chimeras (black bars) imply significant developmental delay. F) Expression of *T* for Limb mesoderm lineage. The line is a curve showing smoothed expression (Loess curve). G) perturbSuite_DE_static applied to chimera Lateral Plate Mesoderm (LPM) cells along the Limb mesoderm lineage identifies genes that are differentially expressed with respect to internal and external control. H) Combining perturbSuite and MAYA reveals pathway deregulation for *T^-/-^* knockout cells at the LPM stage of the limb mesoderm trajectory.

Second, at the lineage level, results from analysis with perturbSuite_DA_lineage were broadly concordant with the cell type level results, with *T*^-/-^ cells significantly depleted in trajectories towards Notochord, Somitic Mesoderm, Presomitic Mesoderm, Allantois and Mesenchyme (Fig. 2C, Supp. Fig. 2C).

Third, among the trajectories containing sufficient cells to assess lineage progression in both mutant and WT fraction (Supp. Fig. 2B, 5), perturbSuite_kinetics revealed a delay for both Mesenchyme and Allantois lineage development as a result of *T* knockout (Fig. 2DE). We also observed a delay in the trajectory leading to an ectodermal cell type, the Ventral Forebrain Progenitors. Interestingly, our analysis shows a depletion of the Mesenchyme as well as a delay in its lineage development (Fig. 2DE). A role for *T* in cancers derived from mesenchymal tissues has been identified previously, where its overexpression is associated with changes in the epithelial-to-mesenchymal transition [8–10].

As previously noted, *T* knockdown severely disrupts the formation of posterior somitic tissues, see Figure 2A-C. Consistently, Allantois cells, which arise from a posterior mesodermal precursor population, are also significantly depleted in *T* chimeras, and tdTom^+^ *T*^-/-^ cells are delayed in their progression along the inferred allantois trajectory (Figure 2DE). These results align with well known allantois defects that have been documented for *T* knockout mice and chimeric embryos [11], giving confidence in the performance of perturbSuite.

Noting that *T* is expressed in the Lateral Plate Mesoderm (LPM) at early stages of the limb mesoderm trajectory (Fig. 2F), we used perturbSuite_DE to gain new insights into limb development. Specifically, we noted a significant reduction in *T*^-/-^ cells for the LPM lineage. These findings are consistent with a previous study that showed impaired forelimb bud formation in *T* mutant embryos [12], but the underlying *T*-dependent mechanisms that disrupt forelimb formation had remained unclear [13]. Using PerturbSuite, we identified that *Vim* and *Aldh2* are strongly downregulated (Fig. 2G), both compared to the tdTom^−^ host cells in the same embryos as well as the WT control experiment.

The downregulation of *Vim* suggested that *T* regulates EMT along the limb mesoderm trajectory [14], which was supported by pathway analysis using MAYA [15] and perturbSuite_DE_static (Fig. 2H). Pathway analysis also showed increased oxidative stress (Fig. 2H), in agreement with the observed deregulation of *Aldh2* (Fig. 2G), as *Aldh2* deficiency has been shown to increase oxidative stress [16–18]. Furthermore, three related pathways are also significantly downregulated: peroxisome, reactive oxygen species (ROS) and fatty acid metabolism.

A further interesting observation from our perturbSuite_DE analysis was the upregulation of *Apela* in *T*^-/-^ cells (Fig. 2G), which activates PI3K/AKT/mTORC1 signalling [19] (Fig 2H), with loss of *Apela* being shown to cause defects in early mesodermal derivatives, consistent with a previous study [20].

### A characterisation of *Mixl1* chimeras provides insights into epicardium development

To systematically assess the embryo-wide effects of *Mixl1* knockout we generated *Mixl1^-/-^* chimeras and processed the resulting scRNA-seq data using perturbSuite, following the same steps as for the *T* chimeras. *Mixl1*^-/-^ tdTom+ mouse embryonic stem cells were generated using CRISPR/Cas9 to induce frameshift mutations in exon 2 of the *Mixl1* locus, thereby causing an early stop codon and functional inactivation of the homeobox domain (Supp. Fig. 3A). Two independent *Mixl1*^-/-^ tdTom+ clones were injected into wildtype blastocysts to generate embryonic chimeras, and three independent pools of E8.5 *Mixl1^-/-^* chimeras were collected and sorted by flow cytometry based on the tdTom reporter and analysed by scRNA-seq (see Methods).

Using DA_cell_type and DA_lineage in *Mixl1*^-/-^ chimeras we observed a broad effect on lateral plate mesoderm (LPM) and its derivatives: a depletion of cell types and lineages for Cardiomyocytes of both first and second heart fields as well as Epicardium and Erythroid cells, and an increased representation of LPM (Fig. 3A, Supp. Fig. 2D). Collectively therefore, this observations amount to a depletion of splanchnic mesoderm with simultaneous enrichment of somatic mesoderm. Although *Mixl1^-/-^* mutants have been previously shown to have defective cardiac development [21,22], the molecular role of *Mixl1* for cardiac cell type induction had not been determined. *Mixl1* knockout mice are known not to form a hindgut [21]. Consistently, in the *Mixl1^-/-^*chimeras, Hindgut and Midgut were fully depleted at cell type level, while Foregut, Gut tube, Pharyngeal endoderm and Thyroid primordium were all partially, but substantially, depleted (Fig. 3A).

**Fig. 3:**
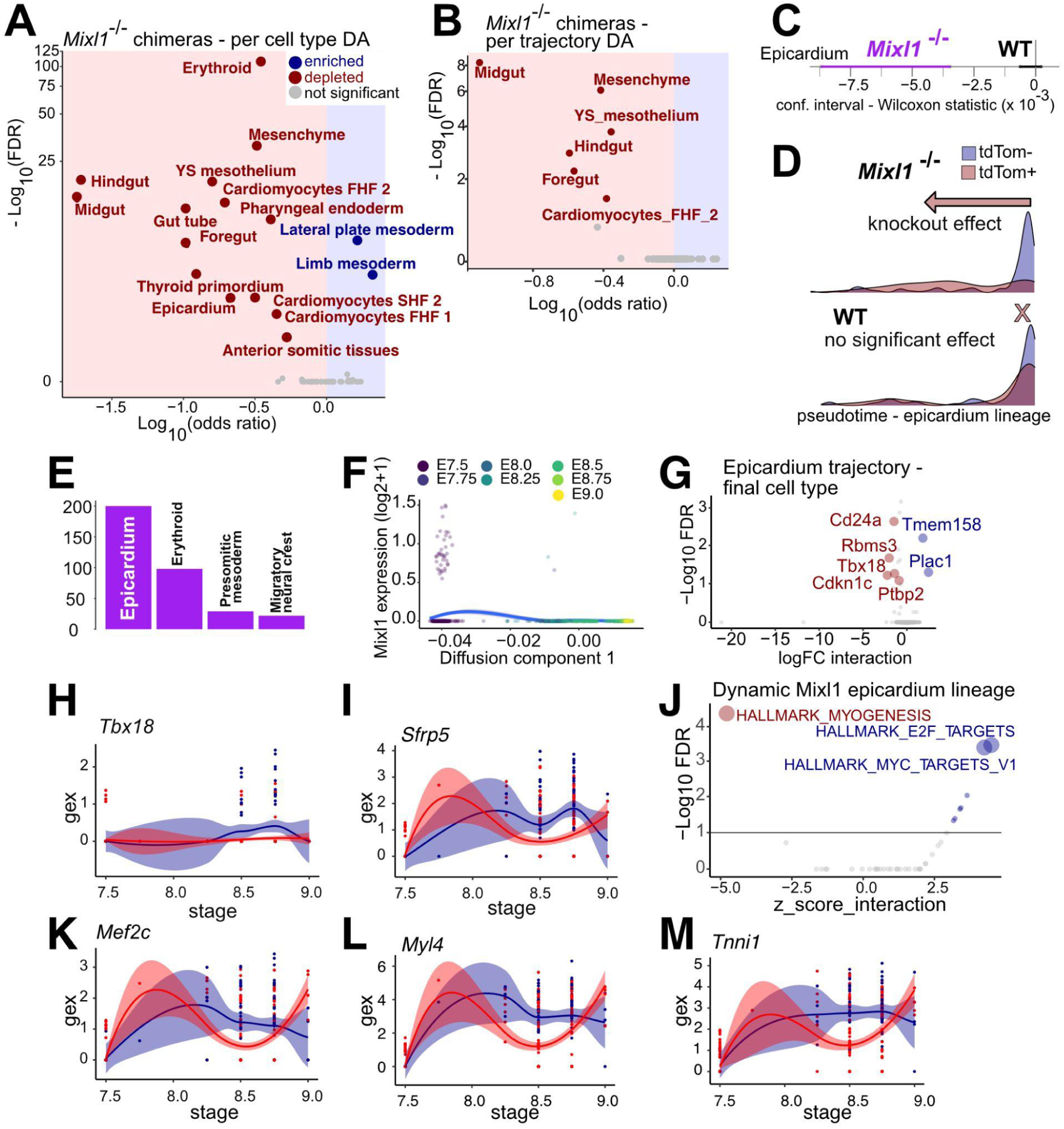
PerturbSuite characterises the effects of *Mixl1* knockout comprehensively and reveals mechanistic insights into Epicardium development. A) DA at the cell type level for *Mixl1*^-/-^chimera knockout cells (perturbSuite_DA_cell_type) B) DA of WOT trajectories leading to the listed cell types at E9.25 (perturb_DA_lineage). C) Confidence interval for location parameter of Wilcoxon test (perturbSuite_kinetics). Confidence intervals for *T*^-/-^ chimeras (purple bars) not overlapping 0 or the confidence intervals for WT chimeras (black bars) imply significance of developmental delay. D) Pseudotime distributions tdTom^+^ (red) and tdTom^−^ cells (blue) within *Mixl1*^-/-^ and WT chimeras. E) Number of genes identified as dynamically DE with PerturbSuite_DE_dynamic, for the four lineages with the most such DE genes. F) Expression of *Mixl1* for Epicardium lineage with smoothed expression (Loess curve). G) PerturbSuite_DE_static for Epicardium state. H-I) Expression levels of *Tbx18* (H) and *Sfrp5* (I). J) Pathways for which *Mixl1* knockout leads to an increase (blue) or decrease (red) in activation along the progression of the lineage. K-M) expression levels of *Mef2c (K), Myl4 (L), Tnni1 (M)*. H, I, K, L, M: The shaded areas correspond to 95% confidence levels for the smoothed expression curve. tdTom+ cells are in red, tdTom- in blue.

After this analysis at the cell type level, we proceeded to investigate lineage development with perturbSuite_DA_lineage. Note that perturbSuite_DA_lineage was not used for Gut tube and Pharyngeal endoderm, as they are below our set frequency threshold at the final time point of E9.25 (Methods, Supp. Fig. 2B, 5B). All of the other cell types mentioned above as depleted at the cell type level were also depleted at the lineage level (Fig. 3B), in line with a general concordance between cell type differential abundance and changes in contribution to corresponding lineage trajectories (Supp. Fig. 2B).

Next, we explored whether there were any alterations in the dynamics of progression along trajectories using perturbSuite_kinetics. Of the depleted cell types with sufficient numbers of cells mapping to a given lineage (Supp. Fig. 2B) only Epicardium showed significantly and substantially altered dynamics (Fig. 3CD). This delay in the Epicardium lineage and the large number of dynamically DE genes (perturbSuite_DE_dynamic, Fig. 3E, Supplemental table 1) suggested an important role for *Mixl1* in epicardium development. Even though *Mixl1* is lowly expressed in the reference atlas and only at early stages of the Epicardium trajectory (Fig. 3F), applying perturbSuite_DE_static revealed significant differences for *Mixl1^-/-^* cells relative to both tdTom^−^ host cells in the same embryos and to the control experiment (Fig. 3G, Fig. 1F). Importantly, we observed downregulation of *Tbx18* (Fig. 3H), a known regulator of the normal structural and functional development of the Epicardium [23]. Our results therefore suggest that expression of *Mixl1* in LPM is required for downstream development of Epicardium and *Tbx18* upregulation.

Applying perturbSuite_DE_dynmaic revealed 199 significantly dynamically DE genes for the Epicardium lineage (Fig. 3E). In particular, we found significant differential dynamics for *Sfrp5* (Fig. 3I), which is expressed in cardiac progenitors, including Epicardium, where it has been suggested to modulate *Wnt* signalling [24].

To better understand the relevance and mechanisms of the dynamically DE genes, we combined perturbSuite_DE_dynamic with pathway analysis (Fig.3J), and identified genes that are members of a de-regulated pathway and that are also identified as dynamically DE themselves. For the strongly downregulated Myogenesis pathway (Fig.3J, Supp. Fig. 3B), *Mef2c* (Fig. 3K), *Myl4* (Fig. 3L), and *Tnni1* (Fig. 3M) had high weight for the pathway score and were also dynamically downregulated on a per-gene basis (Fig 3J, Supp. Fig. 3C). This suggests that in the absence of *Mixl1*, LPM, which gives rise to Epicardium and other cardiac lineages, has a decreased potential towards a cardiomyocyte fate, consistent with the observed depletion of the Cardiomyocytes (Fig. 2A, Supp. Fig. 3D). Furthermore, the Hallmark *E2f* and *Myc* target gene sets are most strongly upregulated (Fig.3J), indicating a persistent dividing progenitor state in mutant cells (that lack the ability to differentiate), consistent with increased abundance of the LPM cell type (Fig. 3A).

### *Mixl1* knockout depletes specific cardiac progenitor populations

Prompted by recent reports suggesting previously unrecognised cardiac progenitor populations [25], we decided to delve deeper into the effects on cardiac development caused by *Mixl1* knockout. In addition to the depletion of cardiac cell types (Fig. 3A-B) and substantial gene dysregulation along the Epicardium trajectory (Fig. 3C-E), we also observed a depletion of cells assigned to the Mesenchyme cell type (Fig. 3AB), a cell type that we found to be part of the Epicardium trajectory (Supp. Fig. 3E). This led us to investigate whether we could identify precursors of the Epicardium within the population labelled as Mesenchyme in the extended mouse gastrulation atlas. Indeed, sub-clustering the Mesenchyme cell type (Fig. 4A) revealed transcriptionally defined subsets, segregated by the expression of markers for a recently identified epicardial and cardiomyocyte progenitor population termed the Juxta-cardiac field (JCF) [25], consistent with the notion that cells annotated as Mesenchyme may comprise JCF cells that would constitute putative progenitors for the cardiac cell types affected in *Mixl1^-/-^*chimeras. Indeed, the average expression of JCF markers (JCF score, JCFS) was downregulated in *Mixl1^-/-^*cells mapping to the Mesenchyme cell type (Fig. 4B, p-value< 2.2e-16 for Wilcoxon rank-sum test), and clusters with high JCFS were strongly depleted in *Mixl1^-/-^*chimeras (Fig. 4C). Altogether this suggests that *Mixl1* functions in heart development include a previously unknown yet critical role in the onset of a JCF program, impacting on the downstream development of cardiac cell types, including the Epicardium.

**Fig. 4:**
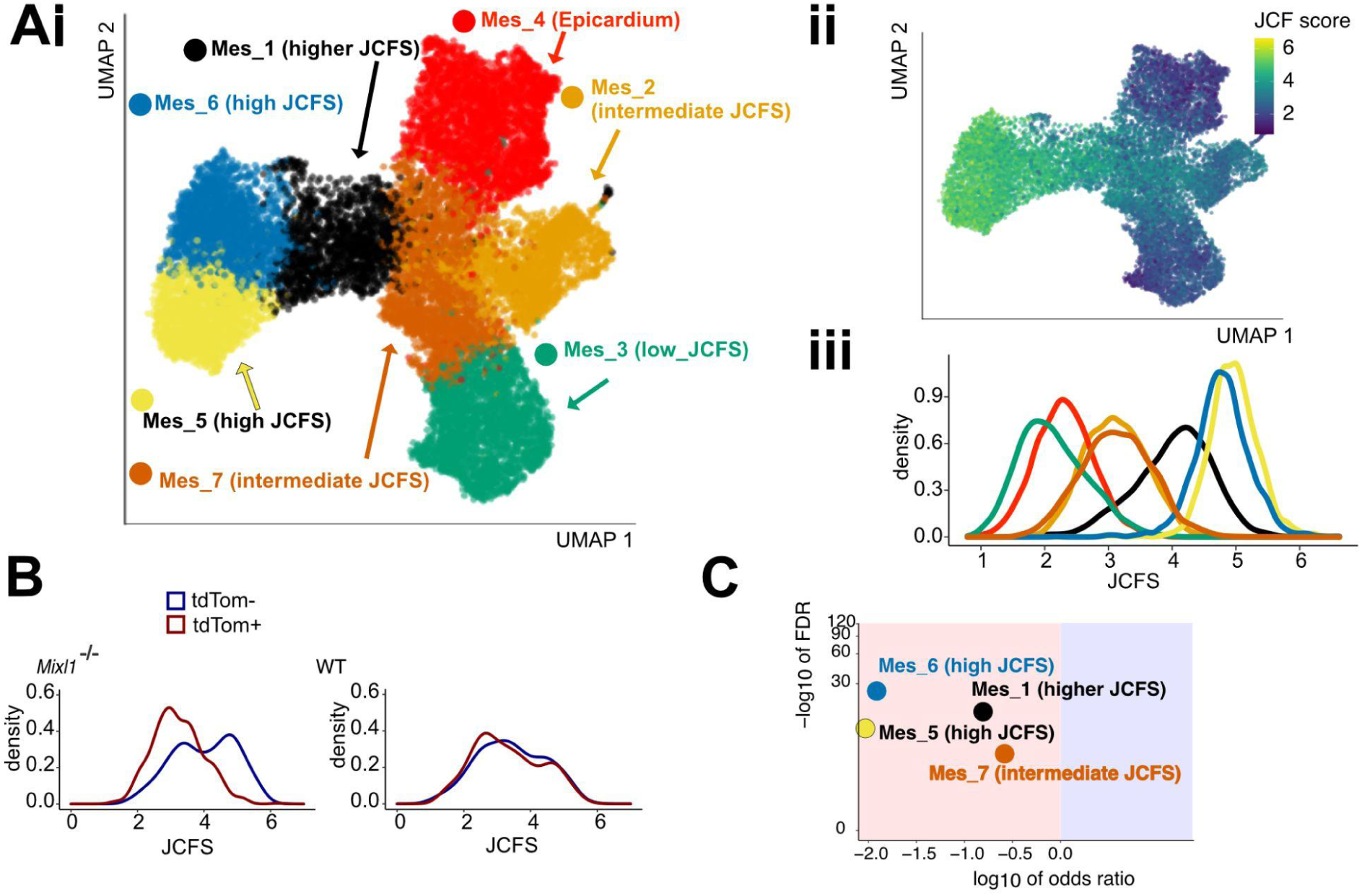
Mesenchyme subclusters into populations with different levels of juxta-cardiac signature. A) Sub-clustering of Mesenchyme. i) Louvain clustering for the extended mouse gastrulation atlas for cells annotated as Epicardium or Mesenchyme based on the whole transcriptome, Uniform Manifold Approximation and Projection (UMAP) coordinates recomputed for Epicardium and Mesenchyme cells only; (ii) UMAP coloured by average log-expression of marker genes of JCF (JCF score (JCFS)); (iii) density plot of JCFS split by cluster show that clusters differ considerably in JCFS levels. Clusters in (i) were labelled according to their levels of JCFS. Cluster Mes_4 mostly comprised the cells labelled as Epicardium in the reference data set. B) Distribution of JCFS for *Mixl1* and WT chimeras across all cells from the Mesenchyme cell type shows a depletion of JCF signature for the *Mixl1^-/-^*cells. C) PerturbSuite_DA_cell_type reveals strong depletion for *Mixl1^-/-^* cells for the clusters with high JCFS. To perform the normalising step (Fig. 1E), we ran perturbSuite_DA_cell_type across all cell types with Mesenchyme replaced by the new subclusters, but only the Mesenchyme subclusters are highlighted in this plot.

### A standardised pipeline streamlines meaningful comparisons

Using perturbSuite, we can easily compare effects of different perturbations if they can be mapped to the same reference framework, in this case the extended mouse gastrulation atlas. As mentioned before, both *T* and *Mixl1* have been reported as major mesoderm regulators [5,21,26,27]. We found that in general, and in particular for a number of specific cell types such as Notochord, Node, Caudal Mesoderm and Caudal Epiblast, *Mixl1* expression was downregulated in *T^-/-^*chimeras (Supp. Fig. 4AB), albeit with small differences in absolute terms, given the generally low expression of *Mixl1* at E8.5. This would suggest a role of *Mixl1* downstream of *T* in the development of these cell types. However, focusing on later cell types we find that these two chimeras generally had distinct phenotypes (Supp. Fig. 4C), and that they also differed largely in terms of the lineages displaying dynamically DE genes (Supp. Fig. 4D).

Interestingly, LPM was depleted in *T^-/-^* chimeras and enriched in *Mixl1^-/-^* chimeras (Supp. Fig4C). Moreover, both chimeras display distinct defects in LPM derivatives, with a depletion of endothelial cells in *T*^-/-^ chimeras compared to defective development of cardiac tissues in *Mixl1*^-/-^ chimeras. This suggests independent roles of the two transcription factors in the development of populations derived from LPM. The population annotated as LPM in the atlas may also comprise heterogenous progenitors with restricted fates, where *Mixl1* and *T* regulate distinct gene networks.

### PerturbSuite can be used to gain insights into AML progression

In addition to the study of the developing embryo, single-cell analysis and transcriptional trajectories may also provide insight into the molecular mechanisms that underpin other processes, such as cancer progression. To illustrate the potential of perturbSuite in this context we applied it to data generated as part of a single-cell RNA-seq study of AML [28]. In this study, the authors used single-cell genotyping along with a machine learning classifier to discriminate mutated from WT cells within individual tumour samples.

We used perturbSuite to gain insight into changes in myeloid development of malignant cells as a result of treatment. To explore this data set within the framework developed originally for chimera analysis as discussed in previous sections, we consider the patient data from day x after diagnosis as corresponding to the knockout chimeras (*T^-/-^* or *Mixl1^-/-^* chimeras), while the D0 data correspond to the WT chimeras. Patient cells classified as malignant correspond to the tdTom^+^ cells, while patient cells classified as WT correspond to tdTom^−^ (Supp. Fig. 6).

To apply perturbSuite, we used the samples from healthy bone-marrow as a reference in lieu of the extended mouse gastrulation atlas for the chimera data sets. We identified genes that are associated with development along the myeloid trajectory, but not with differences between donors (myeloid-trajectory genes). To examine progression, we took advantage of the annotated cell types of this reference data set, which correspond to particular stages along the myeloid trajectory. We then calculated diffusion maps, where annotated cell types follow the expected progression along component 1, from the hematopoietic stem cell (HSC) to the mature cell type (Monocyte) (Fig. 5A).

**Figure 5:**
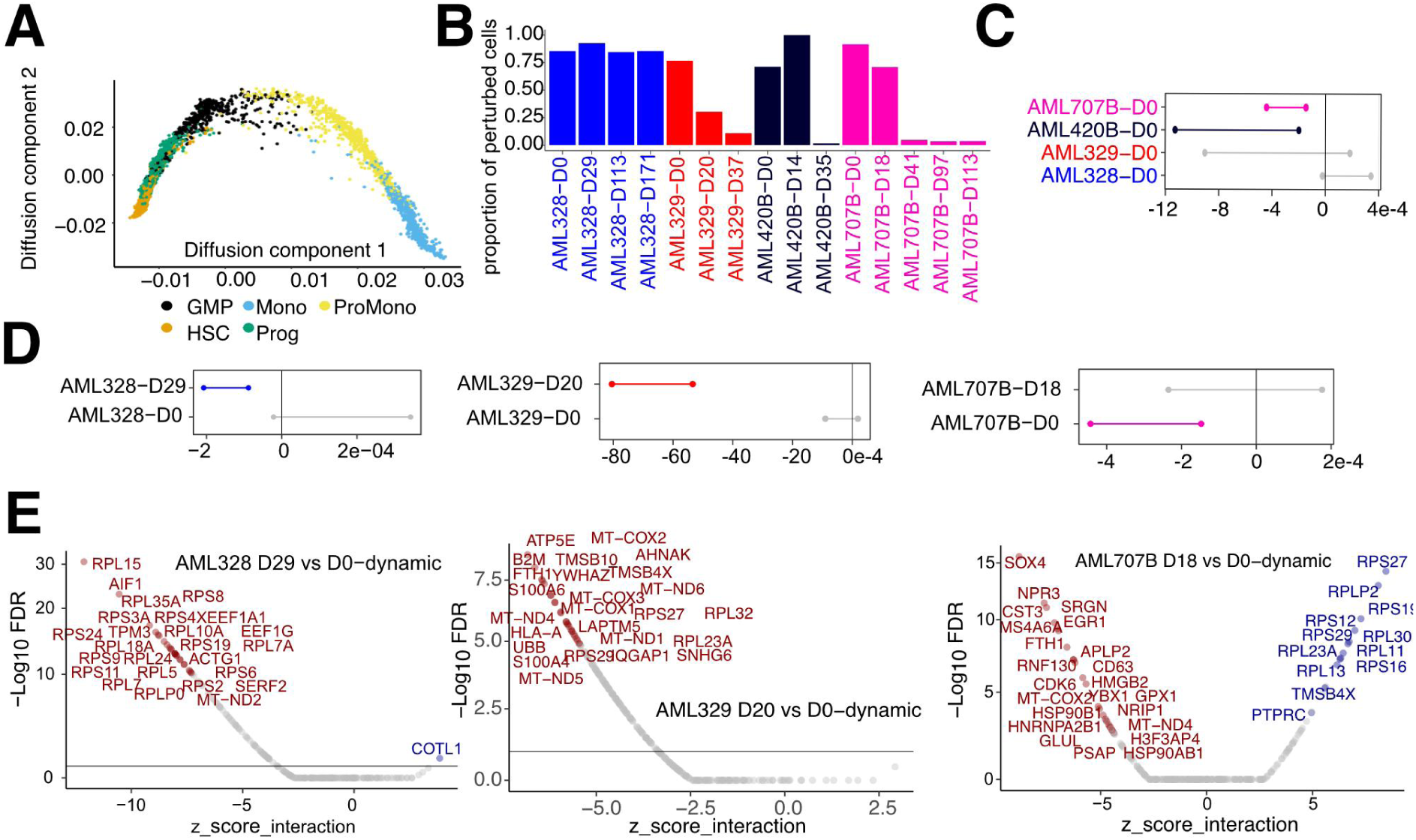
Insights into progression of AML. A) Diffusion components for myeloid trajectory in reference single-cell data set (bone marrow samples from healthy donors), coloured by cell type, as annotated in [28]. B) Proportion of malignant cells along the myeloid trajectory for the four patients with proportions of malignant cells at D0 (diagnosis) less than 95%. C) Wilcoxon test comparing pseudotime along the myeloid trajectory between malignant and normal cells shows a significant delay for patients AML-707B and AML-420B (perturbSuite_kinetics, 95% confidence intervals (bars) below and not crossing 0). D) PerturbSuite_kinetics identifies changes in the delay or acceleration along the trajectory between malignant and normal cells for D0 versus the first time point after diagnosis. AML-420B was excluded from this analysis, as less than 5% of the cells were normal at the first time point after diagnosis. E) Differences in dynamic gene expression changes for the first time point after diagnosis compared to D0, restricted to myeloid-trajectory genes..

Next, we applied perturbSuite_kinetics and perturbSuite_DE_dynamic to investigate how patient samples differ from each other at the time of diagnosis (D0) and in terms of their changes after diagnosis. For this, we focused on patient samples that include multiple time points and for which at D0 less than 95% of the cells along the myeloid trajectory were classified as malignant (Fig. 5B). This ensured a sufficient cell number for both malignant and normal cells at D0 and after treatment, allowing enough power for comparison. We applied perturbSuite_kinetics to test for delay along the myeloid trajectory for malignant cells compared to their WT counterparts. At D0, AML420B and AML707B both show a significant delay, which means that malignant cells are more stem cell-like (Fig. 5C).

While van Galen et al. [28] already showed that the presence of a specific cell type composition in progenitors is strongly related to worse prognosis (Fig. 4ef in van Galen et al. [28]), perturbSuite offered a more comprehensive picture. perturbSuite_kinetics allowed us to study the differences between the location on the myeloid trajectory between malignant and normal cells of the same patient and to follow the composition over time and test for significance of changes over time. We examined in this way the 3 patients from Fig. 5B for whom the percentage of malignant cells in the myeloid trajectory is between 5% and 95% at the first time point after D0 (Fig. 5D).

Interestingly, for AML328 where there is no induction therapy after diagnosis, the malignant cells at the first time point after diagnosis were delayed compared to the normal ones, while there was no delay at D0. For AML707B and AML329, where induction therapy was applied after D0, we observed different patterns. In AML707B the initial delay was no longer present at the next time point, while in AML329 the malignant cells showed a further delay following induction therapy. These differences can be attributed to the diverse mutational landscape of these patients [28], different disease stages as well as heterogeneity of response to therapy. As shown in Fig. 5B, the proportion of malignant cells did not decrease for AML328 over the course of time, while it did for AML329 and AML707B.

We then applied perturbSuite_DE to examine dynamic gene expression effects (Fig. 1F) between D0 and the first time point after diagnosis, based on the myeloid-trajectory genes identified above. In both AML328 and AML329, the differentiation delay of mutant cells in the first time-point sampled after day 0 is accompanied by a decrease, along the differentiation trajectory, of ribosomal protein gene expression compared to normal cells in the same patient (Fig. 5E, Supplemental table 2). In fact, for AML328, 41 of the 85 significantly down-regulated myeloid-trajectory genes are part of the KEGG ribosome pathway, while only 22 out of the 179 not significantly downregulated myeloid-trajectory genes overall are part of the pathway (p=8e-10, Fisher’s exact test). For AML329, 54 of the 140 down-regulated genes are part of the ribosome pathway, compared to 9 out of the 124 not downregulated genes (p=10e-10, Fisher’s exact test). Interestingly we observe the opposite pattern in AML 707B, where the initial delay is no longer present after induction therapy. Here ribosomal protein gene expression increases along differentiation in mutant cells (12 out of 16 upregulated genes are ribosomal, 51 of 248 not upregulated genes, p=1e-5). Of note, these cells also show dysregulation of genes previously implicated in AML development, namely *SOX4* [29], *PTPRC*/CD45 [30] and *EGR1* [31].

## Discussion

With the increasing number of large-scale perturbation experiments [32] in complex systems in *vivo* [2,33] and in *vitro* [3], as well as the emergence of single-cell analysis to patient material [34], principled analysis of large-scale complex perturbation data will allow the identification of lineages of interest, leading to new biological and clinical insights.

Single-cell readouts for perturbation experiments in complex systems allow insights into molecular, cellular and tissue-scale processes across a wide range of cell types and lineages. Here we used scRNA-seq data generated from chimeras to study the role of the transcription factors *Mixl1* and *T* across all lineages in the developing mouse embryo. For this purpose, we developed a set of computational tools, perturbSuite, which enabled us to tackle the particular challenges of complex perturbation data. Our approach allowed the novel identification of affected lineages, such as Limb mesoderm for *T* and *Mixl1*, and Epicardium for *Mixl1*. We also demonstrated the wider relevance of the proposed analysis methods to general perturbation and disease scRNA-seq data by applying it to a study of AML patient data [28].

We used mutual nearest neighbours [35] to map chimera data to the reference data set, for consistency with the batch correction previously applied to our reference set [7]. However, perturbSuite may be used with any mapping [36–38], cell lineage estimation [39], and pseudotime/cell ordering method. PerturbSuite could also be combined with the transcriptional rank of the individual embryo in their reference data set to which the chimera cell is most similar, a concept introduced in [6], instead of pseudotime.

PerturbSuite is a cell-specific coordinate-based approach, locating each individual cell within a framework of precomputed scores on a reference data set (e.g. the extended mouse gastrulation data set). This includes lineage scores (e.g. WOT) and lower-dimensional representations (e.g. diffusion maps). This coordinate-based approach, where each cell is associated with its own coordinates, differs from neighbourhood-based approaches that look for similar groups (neighbourhoods) of cells between a query and a reference data set[40,41]. This cell-specific coordinate-based strategy maintains the high granularity of the single-cell approach and provides a location of the cell in terms of normal development of a reference atlas, along a specific direction (e.g., lineage).

Implementation of perturbSuite on the *T*^-/-^ chimera data set validated our method by reproducing results from previous analyses [5]. Thanks to increased statistical power due to the extended reference Atlas, we detected additional significant effects of *T* knockout on Notochord (Fig. 2A), strongly supported by the literature [11,26,27].

We then applied PerturbSuite to a newly generated chimera data set targeting another key mesoderm regulator, *Mixl1*. In addition to the expected strong depletion of definitive endoderm tissues [21], we note a marked impact on cardiac lineages, with the Epicardium cell type showing the most dysregulated gene expression. *Mixl1*-KO mice show severe cardiac malformations [22], and Epicardium plays an important role in coordinating the development of the other heart tissues, namely through modulation of the *Wnt* pathway [24]. Our results suggest this could be regulated by *Mixl1* through activation of *Sfrp5*, which is depleted in the epicardium lineage for *Mixl1*^-/-^ chimera cells.

*Mixl1* has been recently implicated in an LPM inducing regulatory transcription factor network conserved in chordates [42], but based on the studied enhancer region this did not extend to mammals. Our observations of depleted splanchnic mesoderm (cardiac tissues) and enrichment of somatic mesoderm (limb) suggests that *Mixl1* may also have a major role in balancing lineage propensity within LPM precursors in the mouse.

Our systematic analysis also allowed us to refine cell typing in the reference data set. We found that the general impact of *Mixl1* knockout on cardiac development is reflected by the failure of *Mixl1* knockout cells to develop a JCF signature. This observation has led to identification of cell clusters with a high JCF signature within the Mesenchyme cell type in the extended mouse gastrulation atlas.

Methods to analyse perturbations in complex systems will be essential if we want to harness the full power of single cell genomics for oncology research, where oncogenic mutations commonly lead to molecular perturbations in a minority population within a complex cellular system. When it comes to therapeutic intervention, strategies need to consider the different responses in perturbed cancer cells vs the remaining healthy populations. In the present study we extend the application of perturbSuite to a published AML data set [28] and characterised individual patient samples in terms of differentiation blockage of leukemia cells and response to treatment. We observed that delayed differentiation was associated with a decreased expression of protein synthesis machinery along the myeloid trajectory. This is particularly relevant as translation control is emerging as a potential therapeutic target for AML [43–45]. More broadly, this extension of the work demonstrated the wider applicability of perturbSuite as illustrated with an application to leukemia patient samples.

## Methods

### Embryo Chimera data

#### T^-/-^ and wildtype chimera data

Count data for *T*^-/-^ and wildtype chimeras was obtained from a previous study [5], via the *MouseGastrulationData* Bioconductor package [46].

#### Reference data

Processed data including dimensionality reduction and Waddington Optimal Transport (WOT) scores for the extended mouse gastrulation data was obtained from Imaz-Rosshandler et al. [7]. Processed data for the earlier mouse gastrulation atlas was obtained via the *MouseGastrulationData* Bioconductor package [46].

#### Mixl1^-/-^ embryo chimera data generation

All procedures involving mouse embryos were performed in strict accordance to the UK Home Office regulations for animal research under the project license number PPL 70/8406.

TdTomato-expressing mouse embryonic stem cells (ESC) were derived as previously described [4]. Briefly, ESC lines were derived from E3.5 blastocysts obtained by crossing a male ROSA26tdTomato (Jax Labs – 007905) with a wildtype C57BL/6 female, expanded under the 2i+LIF conditions [47] and transiently transfected with a Cre-IRES-GFP plasmid [48] using Lipofectamine 3000 Transfection Reagent (ThermoFisher Scientific, #L3000008) according to manufacturer’s instructions. A tdTomato-positive, male, karyotypically normal line, competent for chimera generation as assessed using morula aggregation assay, was selected for targeting *Mixl1*. Two pairs of guides were designed to induce large deletions in Exon 2 of the *Mixl1* locus using the http://crispr.mit.edu tool (guide pair 1: AAGCGGCGCCTTCTGCGAAC and TGCTGGGGCGCGAGAGTCGT; guide pair 2: TTGCGGCGCTGTGGCGCCGA and CGCTCCCGCAAGTGGATGTC) and were cloned into the pX458 plasmid (Addgene, #48138) as previously described [49]. The obtained plasmids were then used to transfect the cells and single transfected clones were expanded and assessed for Cas9-induced mutations. Genomic DNA was isolated by incubating cell pellets in 0.1 mg/ml of Proteinase K (Sigma, #03115828001) in TE buffer at 50°C for 2 hours, followed by 5 min at 99°C. The sequence flanking the guide-targeted sites was amplified from the genomic DNA by polymerase chain reaction (PCR) in a Biometra T3000 Thermocycler (30 sec at 98°C; 30 cycles of 10 sec at 98°C, 20 sec at 58°C, 20 sec at 72°C; and elongation for 7 min at 72°C) using the Phusion High-Fidelity DNA Polymerase (NEB, #M0530S) according to the manufacturer’s instructions. To assess the CRISPR/Cas9 targeting, primers to amplify the targeted regions and including Nextera overhangs were used (target pair 1: F- TCGTCGGCAGCGTCAGATGTGTATAAGAGACAGATTATTCCCGCGGCGTCT; R- GTCTCGTGGGCTCGGAGATGTGTATAAGAGACAGCTCCGAGCTGAACGACGT; target pair 2: F- TCGTCGGCAGCGTCAGATGTGTATAAGAGACAGAGCAGCTCCAGTTCGCAGA and R- GTCTCGTGGGCTCGGAGATGTGTATAAGAGACAGCCCAGTTTGCAGTCTAGAACC), allowing library preparation with the Nextera XT Kit (Illumina, #15052163), and sequencing was performed using the Illumina MiSeq system according to manufacturer’s instructions. One ESC clone from each guide pair transfection showing a frameshift mutation resulting in the functional inactivation of the *Mixl1* locus was selected for injection into C57BL/6 E3.5 blastocysts (Supp. Fig. 3A). Chimaeric embryos were harvested at E8.5, dissected, and single-cell suspensions were generated by TrypLE Express dissociation reagent (Thermo Fisher Scientific) incubation for 7-10 minutes at 37°C under agitation. Single-cell suspensions were sorted into tdTom+ and tdTom- samples using a BD Influx sorter with DAPI at 1μg/ml (Sigma) as a viability stain for subsequent 10X scRNA-seq library preparation (version 3 chemistry), and sequencing using the Illumina Novaseq6000 platform in one full S1 flowcell. In total, three independent pools of chimeric embryos were processed, two pools generated with a clone successfully targeted with guide pair 1 (containing respectively 3 and 6 embryos), and one clone successfully targeted with guide pair 2 (comprising 4 embryos). This resulted in 17393 tdTom^−^ and 18754 tdTom^+^ cells that passed quality control (see “Data processing, QC and mapping to extended mouse gastrulation atlas” below). To exclude transcriptional effects intrinsic to the chimera assay, chimaeric embryos were generated by injecting the parental tdTom^+^ *Mixl1*^+/+^ (WT) line into C57BL/6 E3.5 blastocysts and processed as for the *Mixl1*^-/-^ samples.

### AML data

Annotated AML data [28] was obtained through GEO: GSE116256.

### Computational methods

#### Data processing, QC and mapping to extended mouse gastrulation atlas

For the *Mixl1^-/-^* chimeras, raw data were processed with Cell Ranger without mapping intronic reads, with reads mapped to the mm10 genome and counted with GRCm38.92 annotation to ensure consistency with the reference atlas [7] and the existing *T^-/-^*chimera data set [5]. For the *T^-/-^* chimera we used processed reads [5], for the *Mixl1*^-/-^ we followed the quality control steps detailed in Guibentif et al. [5] and https://github.com/MarioniLab/EmbryoTimecourse2018.

#### PerturbSuite overview

PerturbSuite provides a comprehensive picture of knockout effects in complex organisms at four levels (Fig. 1D). First, we investigated whether knockout causes differential abundance at the level of cell types, compared to changes seen between tdTom^+^ and tdTom^−^ cells from the WT chimeras (perturbSuite_DA_cell_type, Fig. 1F, 2A, 3A). Second, for cell types represented at E9.25, we identified the cells comprising the inferred developmental trajectories ending in the respective cell types (henceforth called lineage), and our method determines whether a knockout will lead to abundance or depletion of cells for the lineage, again compared to changes in the WT chimeras (perturbSuite_DA_lineage, Fig2C, 3B). Third, for lineages that are not severely depleted for knockout cells, our suite of methods also determines whether the progression along the lineage is delayed or accelerated (perturbSuite_kinetics, Fig. 1G, Fig 2DE, 3CD, 5CD). Finally, we determine for which lineages knockout cells are affected by differences in gene expression (perturbSuite_DE, Fig 1H, 2G, 3E-M, 5E, Supp. Fig. 3D, Supp. Fig. 4D), and identify trajectory-specific affected genes featuring putative molecular mechanisms downstream of the knockout.

#### perturbSuite_DA_cell_type: Cell type-based DA testing

Cell type-based DA testing determines whether there is significant depletion or enrichment for tdTom^+^ cells compared to tdTom^−^ in the knockout chimeras, compared to the difference between tdTom^+^ and tdTom^−^ in the WT chimeras (Fig. 1F).

Cell type-based differential abundance testing was performed as follows:

First, we discarded cell types with less than 30 WT tdTom^+^ cells assigned to them. For each other cell type c the following ratios were computed:

r_c,target_ = number of tdTom^+^ cells in knockout chimeras mapping to c divided by the number of tdTom^−^ cells in knockout chimera mapping to c

r_c,control_ = number of tdTom^+^ cells in WT chimeras mapping to c divided by the number of tdTom^−^ cells in WT injected chimera mapping to c

Then we performed normalisation to account for the sampling biases (Fig. 1E) as follows: First we computed medians across all cell types: m_target_ = median(r_c,target_) and m_control_ = median(r_c,control_).

From the knockout chimera we sampled (without replacement) n_target_ cells among the tdTom+ cells and n_target_ x m_target_ cells among the tdTom- cells, where n_target_ = min(number of tdTom^+^ cells in knockout chimera mapping to the cell type, number of tdTom^−^ cells in knockout chimera mapping to the cell type/m_target_). Under the assumption that most cell types are not affected by depletion or enrichment in consequence of the knockout (and that therefore the median ration m_target_ is not affected), this normalisation accounts for the fact that given the depletion of larger cell types, an approximately equal number of tdTom^+^ and tdTom^−^ cells overall will lead to spurious enrichment results for other cell types, if not corrected. We proceeded similarly for the WT injected control chimeras.

After sampling, for each cell type we performed a Fisher’s exact test to determine the odds ratio and p-value of a chimera cell mapping to a particular cell type being significantly more or significantly less likely to be tdTom^+^ than tdTom^−^. We repeated the sampling and the Fisher test 100 times to avoid dependence of the magnitude and significance of the inferred effect on the randomly chosen sample. For downstream analysis we used the median odds ratio and the median p-value. FDR correction [50] was subsequently applied to the p-values. We applied perturbSuite_DA not only to cell types, but to a combination of cell type and the embryonic stage of a chimera cell (obtained from mapping to the reference atlas, see the section *Data processing, QC and mapping to extended mouse gastrulation atlas*), as described above with cell type+stage combinations replacing cell types.

#### perturbSuite_DA_lineage: lineage-based DA testing

PerturbSuite_DA_lineage tests whether tdTom^+^ chimera cells have an increased or decreased predisposition to develop into a particular cell type at a later stage (Fig 1C).

The test is similar to perturbSuite_DA_cell_type, but needs to be adapted as we now use lineage scores, i.e. probabilities of a cell being part of a lineage, for each cell rather than cell types (definite assignments to a cell type).

We only considered lineages whose final cell types are present with more than 100 cells at E9.25 in the extended gastrulation atlas, and performed 10 iterations of the following:

> For each cell from the chimeric embryos a lineage was sampled according to the WOT scores. Given the sampled lineage for each cell (a fixed state and not a probability of being in a state), we could then apply perturbSuite_DA_cell_type, performing the sampling of cells and Fisher tests were performed 30 times as for perturbSuite_DA_cell_type.

We then computed the medians across odds-ratios and FDR-corrected p-values across the 10 repeats.

#### perturbSuite_kinetics: testing for acceleration or delay of knockout cells along a trajectory

To determine whether there is a delay or acceleration for tdTom^+^ cells along a trajectory, we combined WOT scores with statistical modelling and pseudotime.

Cells were selected as being part of a lineage using a mixture model of skewed t-distributions [51] with two or three components (the optimal number was determined by the Bayesian information criterion) to identify for each mapped stage which cells belong to the lineage. First we preselected those cells that have higher normalised WOT scores than random, i.e. higher than the inverse of the total number of cells at the stage. Then, we fitted the mixture model and selected from among the cells that passed the first threshold of having WOT scores above random those that were in the cluster of the highest WOT scores. For each stage we discarded cell types that did not constitute at least 10% of the cells for the lineage at that stage, to avoid mistakes from the mapping algorithm to influence the trajectory. From the time point onward from which the final cell type is the most frequent cell type in that trajectory we excluded other cell types from the trajectory.

After the assignment of cells to lineages, temporal genes were selected by testing their association with embryonic time. Then a linear regression model was used to obtain residuals for the association of gene expression with embryonic time for these temporal genes. Genes for which the residuals from that regression were associated with batch, i.e. genes correlated with time to different degrees for different batches, were removed from the list of temporal genes. Then we used diffusion maps [52,53] to obtain a more fine-grained resolution of temporal development, identifying pseudotime as the diffusion component most correlated with actual time, to avoid focusing on components not related to time, but to e.g. spatial location. As the starting point for the trajectories we generally used E7.5, as most chimeric tdTom^+^ and tdTom^−^ cells mapped between E7.75 and E9.0 (Supp. Fig. 1D). Starting from E7.5 allowed for enough power resulting from a sufficient number of mapping cells, as well as sensitivity to lineage delay.

This method was applied to all lineages that were not severely depleted for tdTom^+^ chimera cells, i.e. for which the odds ratio of (tdTom^+^ cells for knockout chimera/tdTom^−^ cells for knockout chimeras)/(tdTom^+^ cells for WT chimera/tdTom^−^ cells for WT chimeras) > 0.05. Pseudotimes were assigned to chimera cells by assigning to the chimera cell the average of the pseudotimes of the 10 atlas cells the chimera cell is most correlated to, where for the computation of the correlation expression we only used genes that were identified as temporal genes in the previous step using atlas cells only. This ensured the pseudotime mapping was based on genes that differ across stages in the reference data set and excluded genes whose expression depends on batch.

We used the Wilcoxon rank sum test [54] to identify trajectories whose median was more strongly shifted in tdTom^+^ cells for the knockout chimeras than for the WT chimeras (Fig. 1G, Fig. 2DE, Fig. 3CD, Fig. 5CD). This identified trajectories for which the difference between tdTom^+^ and tdTom^−^ cells for the knockout chimera was signficant and the 95% confidence intervals for the location parameter (the difference in pseudotime median of the distribution of the tdTom^+^ and tdTom^−^ cells) of the Wilcoxon test were non-overlapping between WT and knockout chimeras (Fig. 2E, 3C, Fig. 5CD).

#### perturbSuite_DE_static: lineage-specific static DE test

The idea of testing for significance of differences between tdTom^+^ and tdTom^−^ cells for each lineage was extended to changes not only in pseudotimes between the tdTom^+^ and tdTom^−^ cells (perturbSuite_kinetics), but to differences in gene expression between tdTom^+^ and tdTom^−^ cells specific to the knockout chimeras as opposed to the WT chimera experiment (Fig 1H). To investigate this, for each gene *j* identified as relevant to the development of the specific lineage (based on the atlas cells), the following mixed model was used:

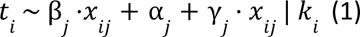

where 𝑡_𝑖_ =1 if cell *i* is tdTom^+^ (𝑡_𝑖_ =0 if cell *i* is tdTom^−^), 𝑥_𝑖𝑗_ is the expression level of gene *j* in cell *i* and and 𝑘_𝑖_ is an indicator of whether the cell is from the knockout (*T* or *Mixl1*) chimera (𝑘_𝑖_ =1) or the WT chimera (𝑘_𝑖_ =0). The parameters β_𝑗_, α_𝑗_ and γ_𝑗_ are inferred. α_𝑗_ is the intercept parameter, the level of gene expression independent of the tdTom status of a cell or its membership in the knockout or the WT chimera. β_𝑗_ corresponds to the added level of gene expression (increase if β_𝑗_ > 0, decrease if β_𝑗_ < 0) resulting from the cell being tdTom^+^ rather than tdTom^−^. Finally, γ_𝑗_ is the value that needs to be added to β_𝑗_ (negative or positive), if the cell is from the knockout rather than the WT chimera. Thus, testing for whether γ_𝑗_ is significantly different from 0 informs about whether the tdTom status (tdTom^+^ versus tdTom^−^) is better explained by a specific parameter for knockout (β_𝑗_ + γ_𝑗_) and WT chimeras (β_𝑗_), rather than by one shared parameter. Thereby we were able to infer whether the relation between the expression of gene *i* and the tomato status was different for the target compared to the WT chimeras.

To ensure that the effect was significant not only compared to the WT chimeras, but also per se for tdTom^+^ versus tdTom^−^ cells of the knockout chimera itself, the following model was used to test for β_𝑗_ ≠ 0, where *i* was now restricted to the knockout chimera cells, i.e. the model is only used for the knockout chimeras.

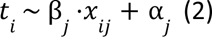

In summary, if γ_𝑗_ ≠ 0 for model (1) and for β_𝑗_ ≠ 0 for model (2) (significantly), then the gene is differentially expressed for tdTom^+^ compared to tdTom^−^ cells for the knockout chimera (β_𝑗_, internal control), and that difference is significantly larger or smaller than the analogous difference for the WT chimera experiment (γ_𝑗_, external control).

To avoid confounding DA of different cell types or lineages along the lineage for tdTom^+^ versus tdTom^−^ cells with DE (and therefore spuriously detect DE of cell type or lineage markers between tdTom^+^ and tdTom^−^ cells), we restricted perturbSuiteDE_static to testing DE for individual cell types (Fig. 2G). We also applied perturbSuite_DE_static to pathway scores inferred using MAYA pathway activity scores [15] instead of individual genes (Fig. 2H).

#### perturbSuite_DE_dynamic: Lineage-specific dynamic DE test

Static DE was then extended to dynamic effects along pseudotime (Fig. 1H), using the following model:

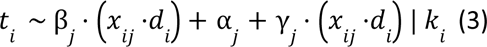

where 𝑡_𝑖_ =1 if cell *i* is tdTom^+^ (𝑡_𝑖_ =0 if cell *i* is tdTom^−^), 𝑥_𝑖𝑗_ is the expression level of gene *j* in cell *i*, *d_i_* is the pseudotime of cell *i*, and and *k_i_* is an indicator of whether the cell is from the knockout chimera (𝑘_𝑖_ = 1) or the WT chimera (𝑘_𝑖_ = 0). The difference between models (1) and (3) is that model (3) tests the change (increase or decrease) of gene expression along the lineage using the interaction between pseudotime and gene expression (𝑥_𝑖𝑗_ ⋅𝑑_𝑖_), rather than test for differences in static gene expression.

Similar to perturbSuite_DE_static, we also checked with perturbSuite_DE_dynamic whether the effect was significant for the knockout chimeras per se, and not only compared to the WT.

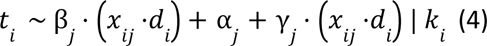

In summary, if γ𝑗 ≠ 0 for model (3) and for β𝑗 ≠ 0 for model (4) (significantly), then the way that the gene changes its expression along the trajectory is different for tdTom^+^ compared to tdTom^−^ cells for the knockout chimera (β_𝑗_, internal control), and that difference is significantly larger or smaller than the analogous difference for the WT chimera experiment (γ_𝑗_, external control).

As perturbSuite_DE_dynamic compares how expression changes along the trajectory, there are, unlike with perturbSuite_DE_static, no confounding effects with differential kinetics along the lineage or DA of cell type along the lineage. Finally, we also combined perturbSuite_DE_dynamic with MAYA pathway scores (Fig. 3J).

#### Sub-clustering of Mesenchyme cell type

We used Louvain clustering [55] on a nearest neighbour graph with k=20 neighbours. To enable detection of genes differentially expressed across sub-clusters without inflation of the false positive rate, we used only 50% of the cells in the clustering algorithm and assigned the remaining cells to the clusters using support vector machines [56].

## Availability of data and materials

Raw sequencing data for the Mixl1 chimeras has been submitted to the Arrayexpress database (http://www.ebi.ac.uk/arrayexpress) under accession number E-MTAB-13409.

Code is available on GitHub (https://github.com/MarioniLab/perturbSuite).

## Acknowledgements

We thank William Mansfield at and the Wellcome-MRC Cambridge Stem Cell Institute animal facility for blastocyst injections, the Flow Cytometry Core Facility at the Cambridge Institute for Medical Research for cell sorting, Katarzyna Kania and the CRUK-CI genomics core for preparing the 10X libraries and for sequencing and Jonathan Griffiths for comments on the manuscript. M.E.S. is supported by the Wellcome Trust (220442/Z/20/Z). C.G. was funded by the Swedish Research Council (2017-06278) and by a Swedish Childhood Cancer Fund position grant (TJ2021-0009). M.-L.N.T. was funded by a Herchel Smith PhD Fellowship in Science. S.M. is funded by a British Heart Foundation PhD Studentship. L.T.G.H. was funded by a Wellcome Early-Career Award (226309/Z/22/Z). R.T. was funded by the British Heart Foundation Cambridge Centre of Research Excellence (RE/18/1/34212) and core support from Wellcome to the Wellcome-MRC Cambridge Stem Cell Institute. Work in the Gottgens group is supported by Wellcome, Blood Cancer UK, MRC and CRUK, and by core support grants from Wellcome to the Wellcome-MRC Cambridge Stem Cell Institute. This work was funded as part of a Wellcome grant (220379/B/20/Z) awarded to B.G. and J.C.M.

## Author information

### Contributions

M.E.S. analysed the *Mixl1* embryonic chimera data set, wrote all code, implemented perturbSuite and applied it to all data sets presented; M.-L.N.T. performed *Mixl1* chimeric embryo dissections and generated the scRNA-seq data set; C.G. generated the *Mixl1* knockout ESC lines; S.M. genotyped *Mixl1* ESC knockout clones; J.B. assisted with embryo dissections; N.W. assisted in scRNAseq data generation; M.E.S, C.G. and J.C.M. interpreted the results with input from L.T.G.H., I.I-R., M.-L.N.T.,R.T. and B.G.; I.I-R. provided processed data and WOT analysis for the extended mouse gastrulation atlas; M.E.S. wrote the first draft of the manuscript; M.E.S., C.G., L.T.G.H., M.-L.N.T., R.T., B.G., and J.C.M edited the manuscript; J.N., B.G., J.C.M. and C.G supervised the study.

## Ethics declarations

### Ethics approval and consent to participate

All animal experiments were performed in strict accordance to the UK Home Office regulations for animal research under the project license number PPL 70/8406.

### Declaration of interests

J.C.M. has been an employee of Genentech since September 2022. The remaining authors declare no competing interests.

## Supplementary Figures

**Supplementary Figure 1:**
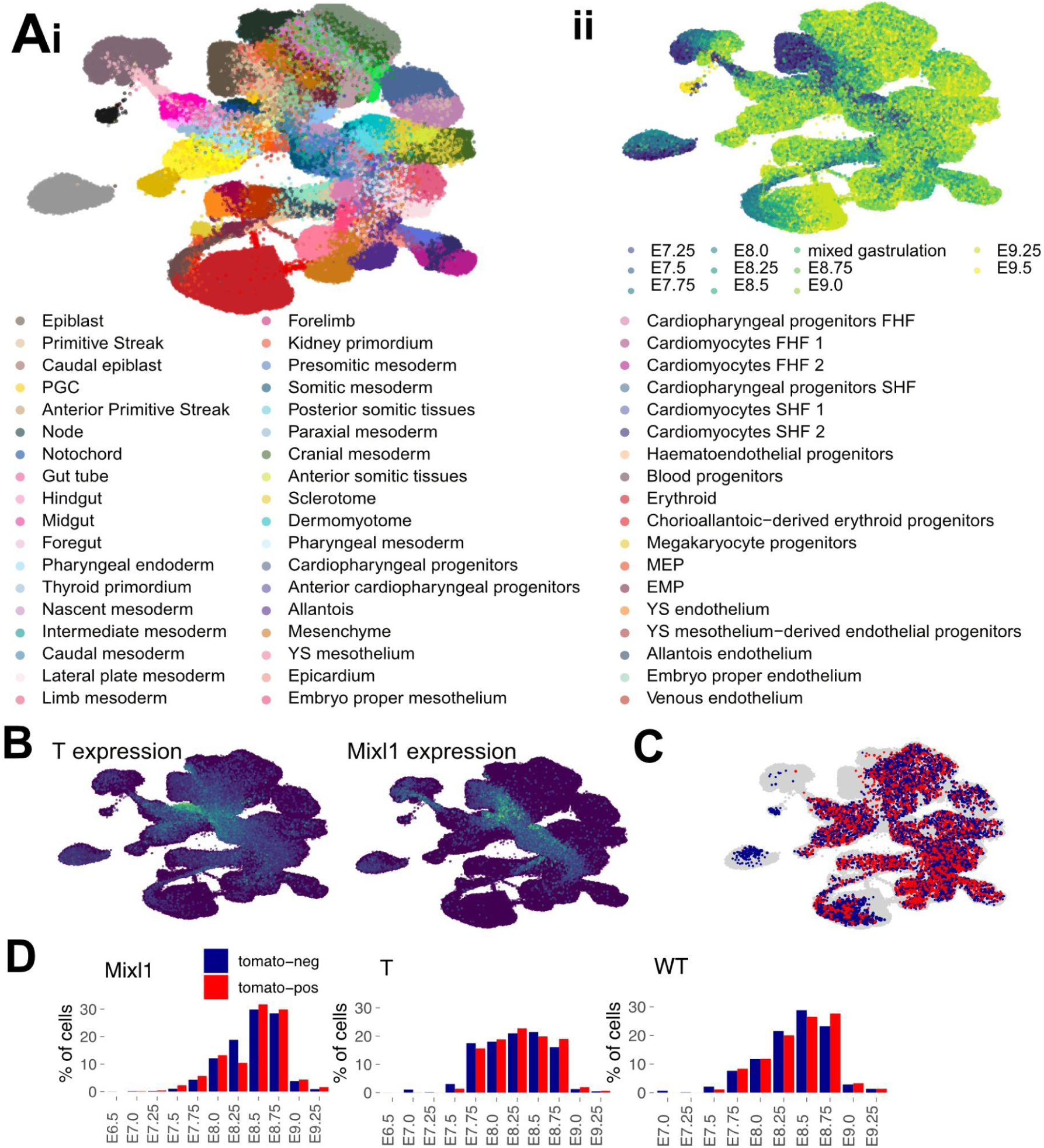
Modelling based on the extended mouse gastrulation atlas. A) UMAP representation of extended mouse gastrulation atlas, coloured by cell types [7] (i) and stages (ii) B) Expression of *T* and *Mixl1* in the extended mouse gastrulation atlas. C) Mapping of the WT chimera to the extended mouse gastrulation data set shows slight differences between injected tdTom^+^ (red) and host tdTom^−^ (blue) cells. D) Mapping of chimeras harvested at E8.5 to different stages in the extended mouse gastrulation atlas. The actual time of collection is E8.5, but as expected we observe differences in developmental progression within the embryo litters. Thus some cells are delayed in development (earlier than E8.5), and others advanced.

**Supplementary Figure 2:**
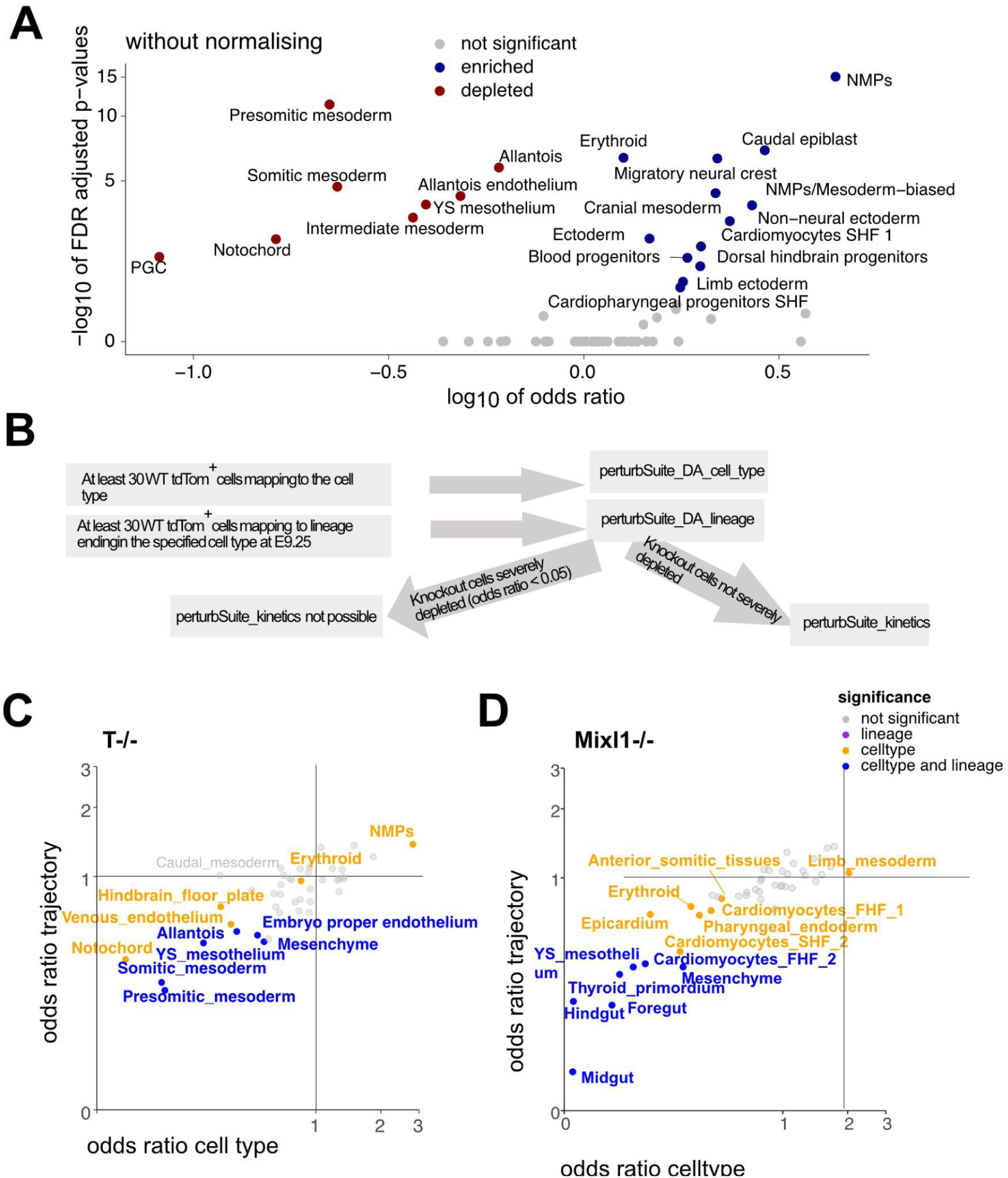
DA analysis. A) DA analysis without normalisation does not account for sampling bias and leads to more cell types appearing as enriched. B) Schematic of applicability of perturbSuite_DA_cell_type, perturbSuite_DA_lineage and perturbSuite_kinetics, which rely on sufficient cells being present in the analysed cell type/lineage. See also Supp. Fig. 5. C and D) Comparing lineage-based to cell type-based enrichment and depletion of *T*^-/-^(C)/*Mixl1*^-/-^(D) cells.

**Supplementary Fig 3:**
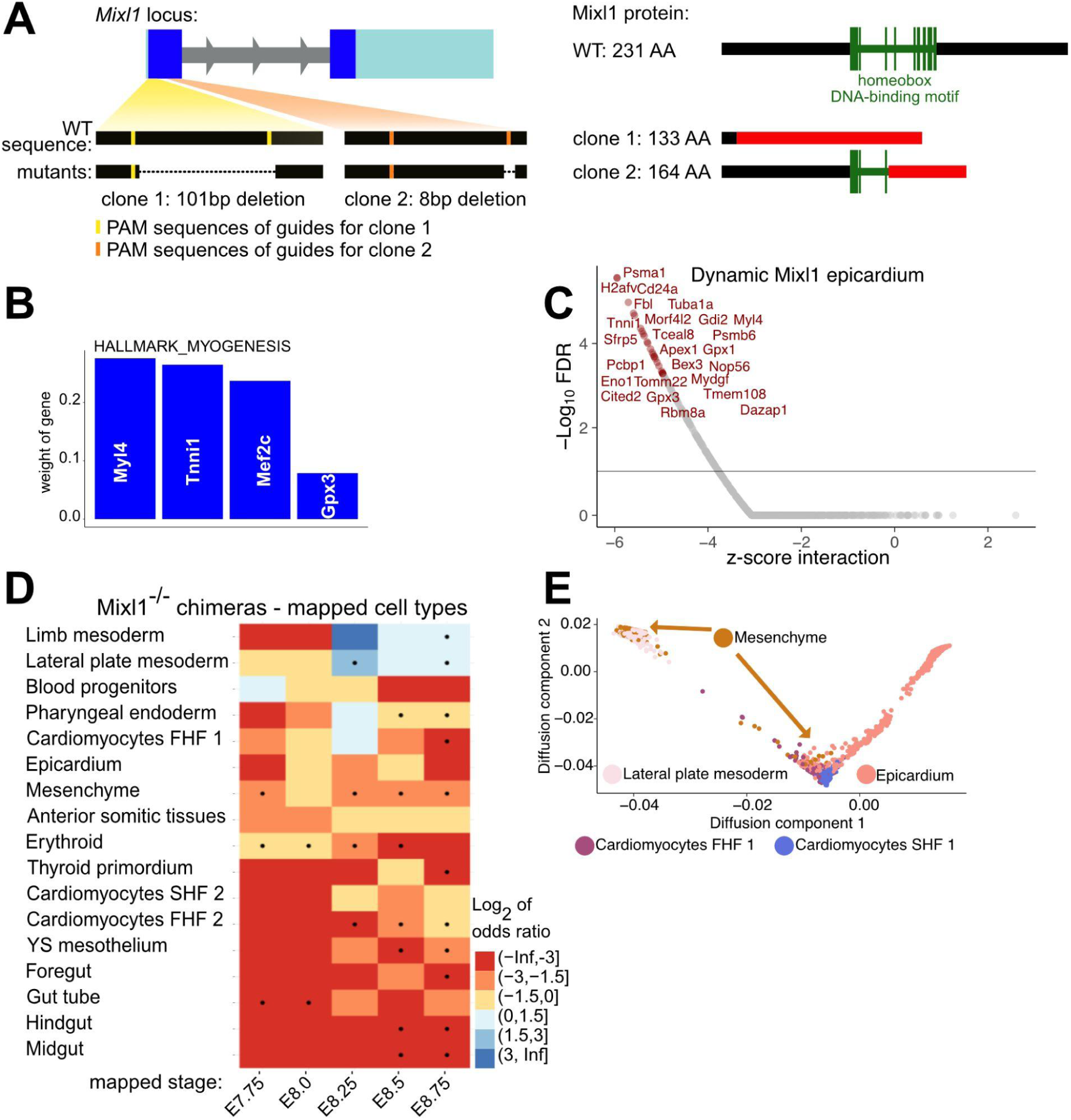
Mixl1^-/-^ chimeras generation and analysis. A) Left: schematic of the *Mixl1* locus, with the CRISPR/Cas9 deletions leading to the generation of each of the *Mixl1* mutant ESC clones. Right: schematic of the *Mixl1* protein, with the effects of the CRISPR/Cas9 deletions on the resulting protein in the mutant ESC clones. Altered protein sequences resulting from the frameshift mutations are depicted in red. B) Weights found by MAYA for the Myogeneis hallmark set. C) Dynamic gene expression changes (perturbSuite_DE_dynamic) in *Mixl1^-/-^* tdTom+ cells for the Epicardium lineage. In red: genes significant for both the internal tdTom^+^ versus tdTom^−^ comparison within the *Mixl1*^-/-^ chimeras, as well as compared to the control experiment. D) DA per mapped stage and cell type for *Mixl1*^-/-^ chimeras. E) Epicardium lineage from E7.5 to E9.0 in the extended mouse gastrulation atlas, coloured by cell type.

**Supplementary Figure 4:**
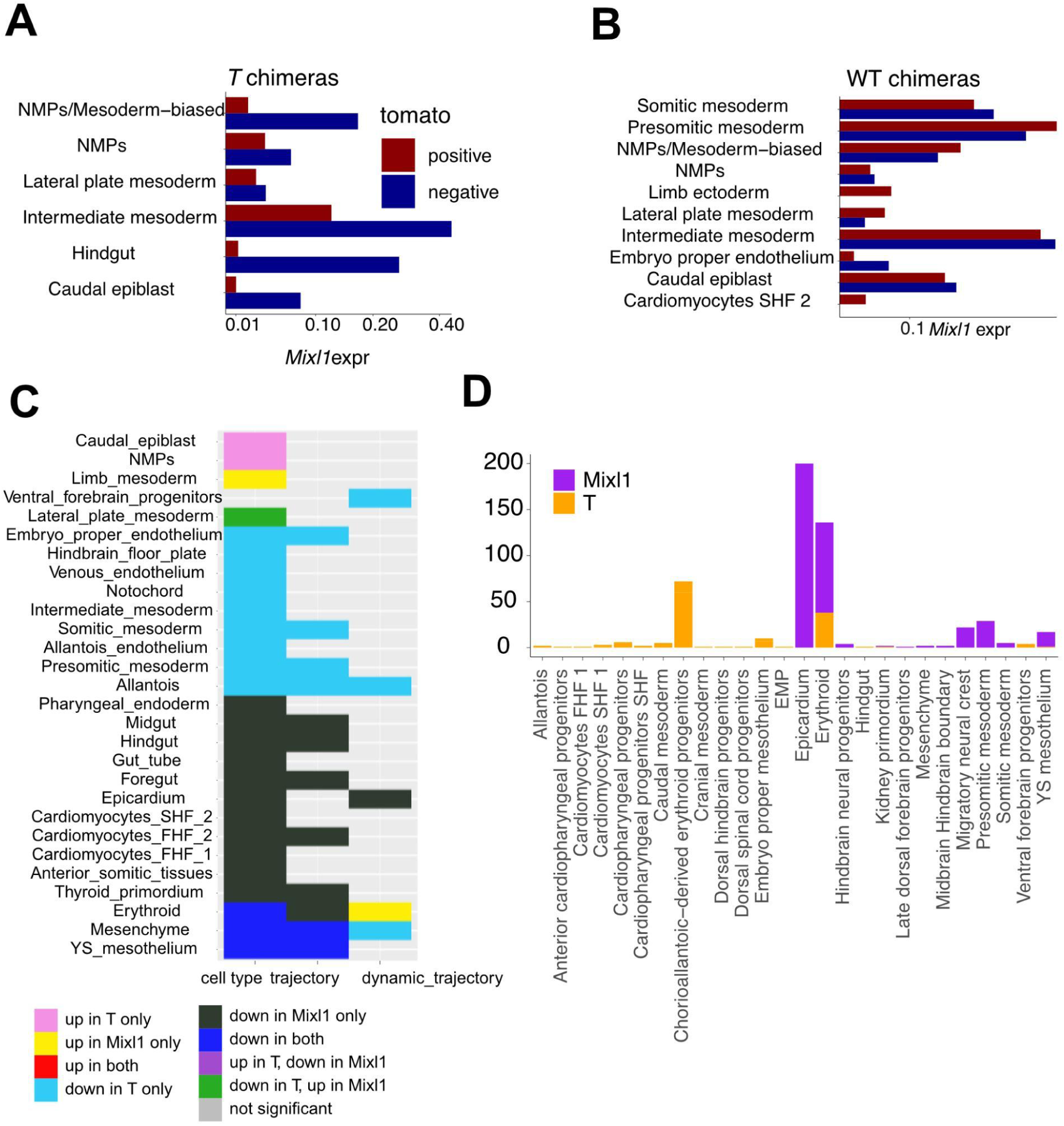
Across-chimera comparison. A) Mean *Mixl1* expression (normalised log scale) in *T*^-/-^ chimeras, for cell types with average normalised log expression of at least 0.033 for either the tdTom^+^ or the tdTom^−^ cells and at least 50 tdTom^+^ cells. B) *Mixl1* expression in WT chimeras, with cut-offs for expression and cell numbers as in A. C) Comparing effects between *T^-/-^* and *Mixl1^-/-^*, for DA of cell type, DA of lineage and differential speed of progression along the trajectory. D) Number of genes identified as dynamically DE for each lineage for *T^-/-^*and *Mixl1^-/-^* chimeras, excluding lineages without any significantly dynamically DE genes.

**Supplementary Figure 5:**
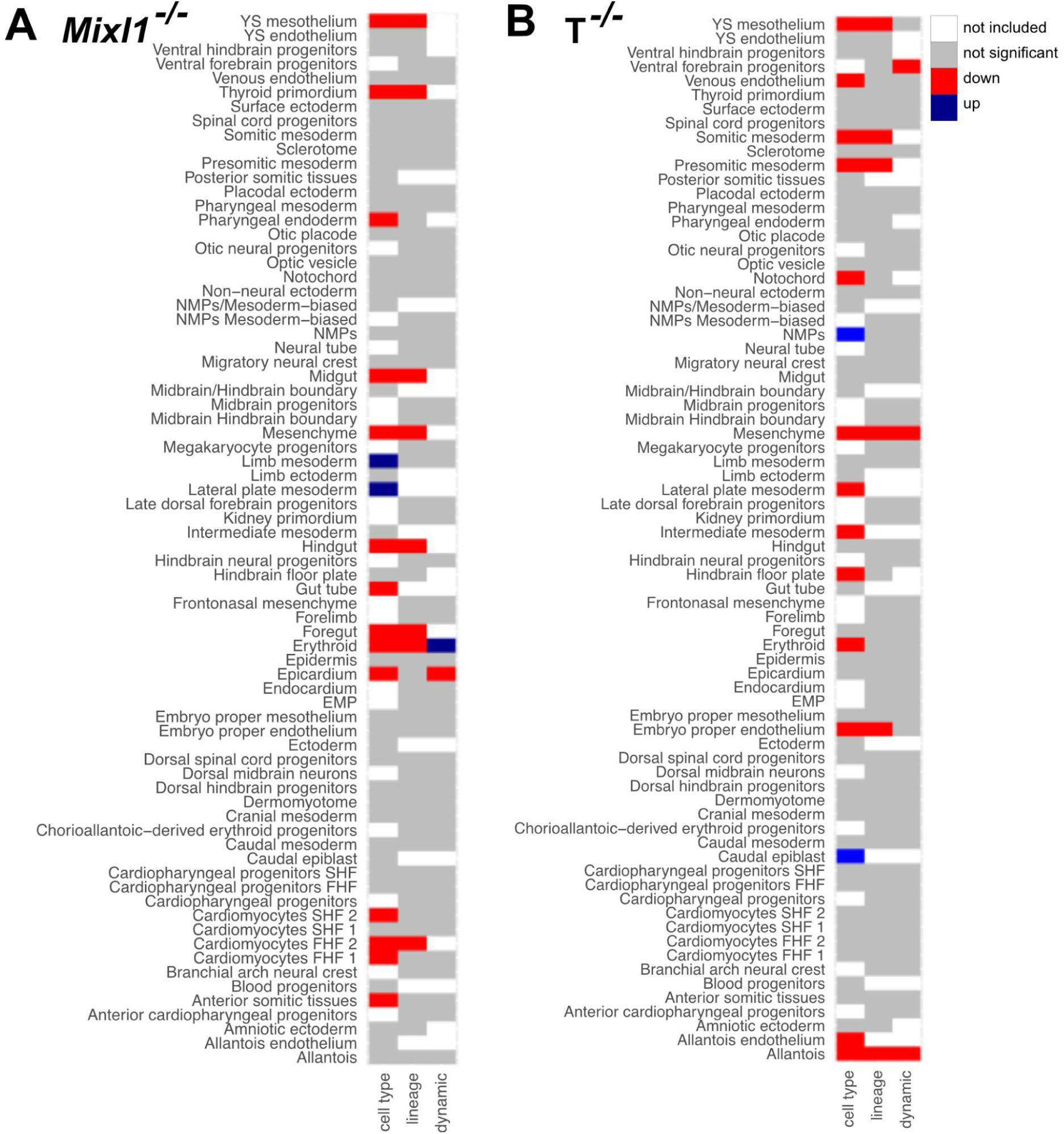
Extension of Supp. Fig. 4c, including all cell types and lineages. Grey colour signifies that a test was performed, but there was no significant enrichment or depletion. White signifies that the test could not be performed due to insufficient representation of tdTom fractions (see Supp. Fig. 2B) for applicability of perturbSuite analyses).

**Supplementary Figure 6:**
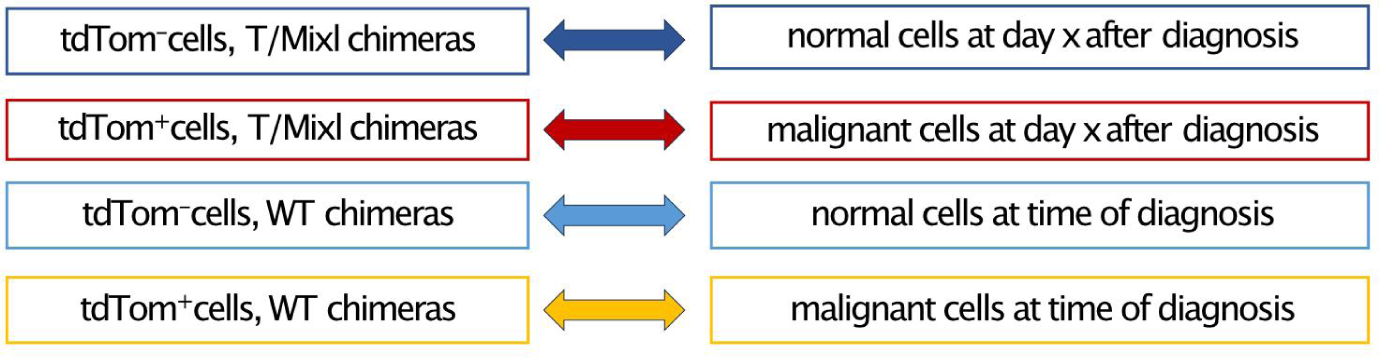
Correspondence between chimera and AML data sets. PerturbSuite_kinetics and perturbSuite_DE_dynamic were applied to the AML data set using internal and external controls similar to the chimera set-up. The figure illustrates which subset of the AML data corresponds to which part of the chimera data.

## Notes

### Summary of Updates

We included new results on heart development and an additional Figure (Fig. 4).

